# Comparative analysis of Pure Hubs and Pure Bottlenecks in Human Protein-protein Interaction Networks

**DOI:** 10.1101/2021.04.06.438602

**Authors:** Chandramohan Nithya, Manjari Kiran, Hampapathalu Adimurthy Nagarajaram

## Abstract

Bottlenecks and hubs form a set of topologically important nodes in a network. In this communication, we have made a detailed investigation on hubs and bottlenecks in human protein-protein interaction networks. We find that, three distinct groups exist which we refer to as: a) pure hubs (PHs, nodes having high degree but low betweenness values), b) mix proteins (MXs, nodes having both high degree and high betweenness values) and c) pure bottlenecks (PBs, nodes having high betweenness values but low degree values). Our investigations have revealed that pure hubs, as compared with MXs and PBs, (i) are more disordered, (ii) have higher potential to bind to multiple partners, (iii) are enriched with essential proteins as well as enriched with a higher number of splice variants. The MX proteins, as compared with PHs and PBs, (i) show slower evolutionary patterns, (ii) are involved in multiple pathways, (iii) enriched with the products of genes associated with various diseases and (iv) are more often targeted by bacteria, viruses, protozoa, and fungi pathogens. PBs, as compared with the PHs and MXs, (i) are associated with cancer genes and (ii) are the targets or the nearest neighbors of the targets of most of the approved drugs. Furthermore, our study revealed that these three categories of proteins are involved in distinct functional roles; PHs are involved in housekeeping processes such as transcription and replication; MXs proteins are involved in core signaling pathways whereas PBs are involved in signal transduction processes. Our work, therefore, has identified the distinct characteristics features associated with pure hubs, mix proteins and pure bottlenecks and thus helps in prioritizing proteins based on their degree and betweenness centrality values.

## Introduction

Proteins are the molecules responsible for many cellular functions that are carried through physical interactions among themselves as well as with other molecules such as lipids, nucleic acid and metabolites. Physical interaction among proteins can be represented as networks or graphs where nodes are proteins and edges are interactions among proteins. Protein-protein interaction networks (PPINs) and their topologies emphasize the functions of proteins^1^, protein evolution^2^ and diseases^3^. Analyses of protein-protein interaction networks (PPINs) have potential to yield detailed information on the system level functionality of a living cell as a function of interacting proteins.

Centrality measures are used for assessing relative importance of proteins in PPINs and correlated to their essentiality in living cells. Among various centrality measures, degree and betweenness are the two fundamental measures in network theory. Degree is defined as the number of interacting partners of a protein while betweenness centrality is computed as the fraction of the total number of shortest paths that pass through a particular node in a network. In a protein-protein interaction (PPIs) network, the nodes with high degree and the nodes with high betweenness have been referred to as hubs and bottlenecks respectively. Hub and bottleneck proteins are topologically important to the network structure. Deletion or removal of hub proteins has been shown to be lethal to the cells.^4^ While the bottleneck protein lies on the communication path and control over information flow between modules in the network which is similar to bridges in highways of transportation networks^5^, disruption of a bottleneck protein could cause partition of the network which is lethal to the cells.^6^

Several studies have investigated various facets of hubs and bottlenecks in PPINs. Hubs have been shown to be significantly enriched with essential proteins in interaction networks,^4,5,7^ whereas bottlenecks are essential in regulatory and other directed networks.^5^ A negative correlation exists between evolutionary rate and degree or betweenness.^8^ The proteins with high degree and also high betweenness are likely to be conserved evolutionary.^9^ It has also been highlighted that hubs and bottlenecks proteins are specially targeted by pathogens.^10,11,12^ Bottleneck proteins are used to identify prostate cancer genes based on network analysis and text mining.^13^ Bottleneck proteins are also involved in neurodegenerative diseases^14^, Breast cancer.^15^ Previous studies have shown relationships between drug targets and betweenness and degree.^16^ The proteins with high degree and betweenness values are suggested to be potential drug targets^17^ and have characteristic properties of successful drug targets. ^18^

In interaction networks, the degree and the betweenness centrality are correlated with each other.^5,19^ As a consequence, a significant overlap exists between hubs and bottlenecks. Considering this aspect, some studies have reported that hub proteins with low betweenness (‘hubs-non-bottleneck proteins’) are more essential than bottleneck proteins with low degree (‘non-hub-bottleneck proteins’).^5^ In addition, it has also been reported that hub-non-bottleneck proteins evolve significantly slower than non-hub-bottleneck proteins.^8^ The non-hub-bottleneck proteins connect functional modules in the PPINs.^9^ Taking cue from these studies we undertook a more detailed and in-depth study on some of the structural/ sequential/functional properties that have not been investigated and could be associated uniquely to the proteins with (a) only high degree values (hereafter referred to as Pure Hubs (PHs)), (b) only high betweenness values (hereafter referred to as Pure Bottleneck (PBs)) and (c) high degree as well as high betweenness values (hereafter referred to as Mix (MXs)) and the details are reported in this communication.

## Materials and Methods

### Protein interaction network

Human protein interactions were collected from Human Integrated Protein-Protein Interaction rEference (HIPPIE) database (Version 2.1, 18 July 2017).^20^ HIPPIE integrates data from experimentally detected interactions databases which consists of MINT,^21^ BioGrid,^22^ DIP,^23^ BIND,^24^ IntAct,^25^ HPRD, ^26^ MIPS. ^27^ HIPPIE confidence score ranges from 0 to 1 which shows the number of experimental studies supporting each interaction. We selected only those interactions with confidence ≥ 0.63 comprising of 2,44,002 physical interactions among 15,251 proteins.

All the protein IDs such as Ensembl, Gene IDs, etc. were converted using a common idmapping file available at “ftp.uniprot.org/pub/databases/uniprot/current_release/knowledgebase/idmapping.dat.gz.”

### Prediction of disordered regions in proteins

The disordered regions in the proteins were identified using a standalone version of IUPred2A.^28^ IUPred2A scores above 0.5 were considered as indicative of disordered state of a residue. The percentage of disordered regions was calculated by counting the number of residues greater than 0.5 and divided by the total protein length. The binding sites in the disordered regions were predicted using ANCHOR2.^29^ The residues with scores > 0.5 were considered as binding site residues.

### Pathway centrality, splice variants and evolutionary rate of the proteins

The pathways were identified by the Kyoto Encyclopedia of Genes and Genomes^30^ (KEGG) annotation data. A total of 337 human pathways were identified from KEGG involving the proteins in the protein-protein interaction network considered in the present work.

The number of splice variants associated with every protein in the network was retrieved from Ensembl^31^ available at ftp://ftp.ensembl.org/pub/release94/fasta/homo_sapiens/pep/Homo_sapiens.Grch38.pep.all.fa.z”.

The evolution rates (dN) of human proteins were calculated with respect to mouse and were obtained from BioMart^32^ (www.ensembl.org/biomart/martview/) (BioMart October 2019)

### Functional enrichment analysis

Gene Ontology (GO) Functional Enrichment analysis was performed using FunRich^33^ standalone tool (V3.1.3) that integrates proteomic and genomic resources. GO analysis was performed for all the proteins that cover the three domains, namely molecular function (MF), biological process (BP) and cellular component (CC) using FunRich database as the background dataset. GO terms with P-values ≤ 0.05 (Benjamini - Hochberg method) were considered significantly enriched in the set of input proteins.

### Identification of proteins from essential genes

We used the Database of Essential Genes^34^ (DEG, V15.2) that contains essential and non-essential genes of bacteria, archaea, and eukaryotic organisms under different environments. This database has information of 8,256 human proteins as on December, 2018. Of these, 7,282 corresponded to the interacting proteins considered in our interaction network.

### Pathogen-Targeted Proteins

Host-Pathogen interaction data were obtained from Pathogen-Host Interaction Search Tool (PHISTO). PHISTO collects experimentally verified protein interactions from STRING,^35^ DIP,^23^APID,^36^ Reactome,^37^ iRefIndex,^38^ MPIDB,^39^ BIND,^24^ MINT^21^ databases. The dataset used by PHISTO contains 26,492 pathogen host interactions between human and 300 pathogen strains (Bacteria, Virus, Fungi and Protozoa). Of these a total of 4,585 human proteins in our protein-protein interaction network are targeted by various pathogens.

### Drug-Target proteins

Data on the drugs and their corresponding human targets were extracted from DrugBank^40^ database (V5.1.4). We could retrieve 1,991 proteins participating in the protein-protein interaction network as the targets of the approved drugs.

### Disease-associated genes

The disease genes were obtained from Online Mendelian Inheritance in Man (OMIM) database,^41^ which is a compendium of human disease genes and phenotype. In OMIM morbid map file, we considered only genetic disorders with symbols “{}” and “(3)”. “{}’’ and their corresponding mutations that are susceptible to infections (eg. Malaria, Diabetes, etc.). As of September 2019, this list contains 5,959 disease genes of which 1,716 corresponded to the interacting proteins in the network considered in this study.

### Cancer -related genes

The cancer related genes were collected from the Genome-wide Association Studies (GWAS) Catalog database (V1.0).^42^ GWAS Catalog contains compilation of human phenotypes and genotypes and their relationships. A total of 2,800 genes given in this catalogue are considered as cancer-associated genes of which 1,247 were found as part of our interaction network.

### Enrichment Score (Z score)

Enrichment of essential gene products, pathogen targeted proteins, drug targets, cancer gene products among the PHs, PBs and MXs was discerned by means of Z-score which is calculated as follows:

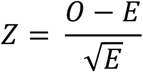

Where, O and E are the observed and the expected number of proteins (PHs, PBs and MXs) in a particular role (essential genes/pathogen targeted/drug targets/cancer genes/disease genes). If Z score is > 1 (O > E), the proteins were considered enriched with the given role whereas proteins with Z score <1 (O < E), were considered depleted in the given role.

### Giant component and path length

The largest connected component in the network is defined as a giant component. The R function “giant.component.extract” was used to extract the giant component from Global Interaction Network (GIN) and this giant component comprises of 15,203 nodes. The characteristic path length is defined as the average shortest path over all the pairs of the nodes in the giant component and this was using igraph of the R package.^43^

## Results and Discussion

### Pure hubs, mix proteins and pure bottlenecks in Human Global Interaction Network

We extracted the known experimentally identified physical interactions among human proteins from the Human Integrated Protein-Protein Interaction rEference (HIPPIE) database.^20^ The resulting GIN comprises of 2,44,002 interactions among 15,251 proteins. The degree and betweenness values for all the proteins in the GIN were calculated using the igraph in R package.^43^ As reported in the literature ^5,19^ the proteins in our network too show a positive correlation between their degrees and betweenness values (Fig. S1).

We considered any protein which is in the top 20 percentile of the proteins in degree distribution (degree ≥ 44) as hub and any protein found within the top 20 percentile in the betweenness distribution (the betweenness values ≥ 14360) as bottleneck. Among these hubs and bottlenecks we could identify three categories: (a) pure hubs (henceforth denoted by PHs) – those proteins with degree ≥ 44 but having betweenness values < 14360), (b) pure bottlenecks (henceforth denoted by PBs) – those proteins with betweenness values ≥ 14360 and degree < 44 and (c) mix (henceforth denoted as MX) – those proteins that hubs as well as bottlenecks. Among 15,251 proteins in the GIN, 926, 629 and 2124 respectively belonged to PHs, PBs and MXs categories (Fig. 1). It is interesting to note that MX proteins are associated with higher degrees and betweenness values as compared with PHs and PBs (Fig.S2).

**Figure 1:**
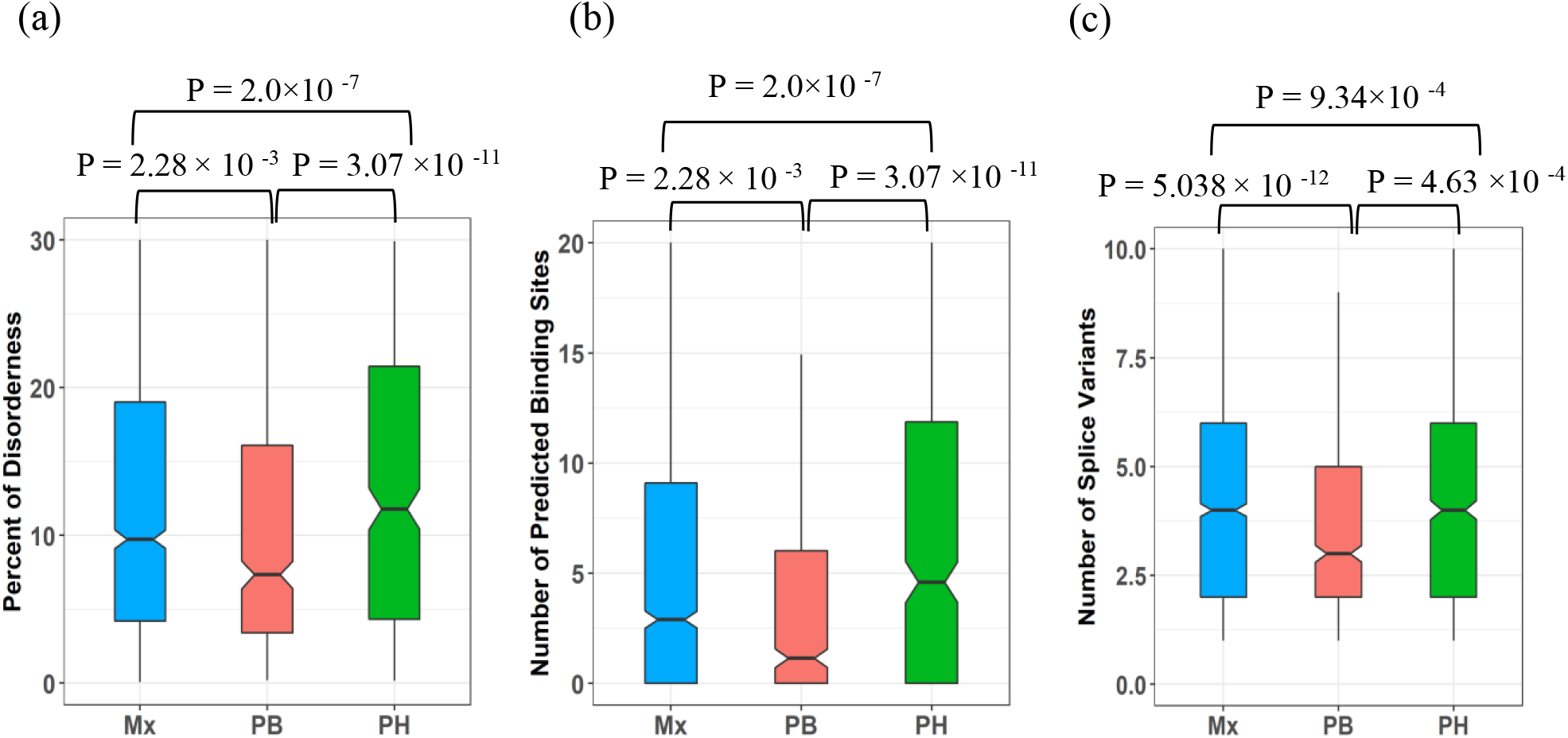
Box plots illustrating (Mx represent mix proteins, PH represent pure hubs and PB represent pure bottlenecks.) (a) Percentage of amino acid residues in the disordered regions, (b) Number of Predicted Binding sites in the disordered regions, (c) Number of Spliced Variants associated with proteins. Pure hubs are more disordered (P value < 2.2 × 10-16) and have more number of predicted binding sites in the disordered regions (P value < 2.85 ×10-11) than pure bottlenecks and mix proteins. While, pure hubs possess more number of splice variants (P value < 4.88 × 10-12) than pure bottlenecks and mix proteins. The P-value was calculated by Kruskal-Wallis test. Outliers have been masked for clarity.

### Disordered regions and predicted binding sites

Intrinsically disordered proteins (IDPs) play an important role in protein–protein interaction.^44,45^ Earlier studies in eukaryotes have shown that hubs are more disordered and have more number of predicted sites in disordered regions.^46,47^ than non-hubs. We compared the disorderedness associated with pure hubs, mix proteins and pure bottlenecks. Among the three categories, we find PHs to be significantly enriched with disordered regions (P value < 2.2 × 10 ^-16^) (Fig.1a) and also have more number of predicted binding sites in the disordered regions of the proteins (P value < 2.85 ×10^-11^) than the other two categories (Fig.1b). The high disorderedness of pure hubs provides flexibility and promiscuity to bind multiple partners at the same or different intervals of time.

### Abundance of Splice variants

In the eukaryotic protein interaction network, a node is not a single protein but an ensemble of its isoforms due to alternative splicing.^48^ It has been reported that hubs are generally associated with a large number of splice isoforms. ^48,49^ We compared the number of splice variants associated with pure hubs, mix proteins and pure bottlenecks. We found that PHs, and also the MXs, are associated with more number of splice variants than PBs (P value < 4.88 × 10 ^-12^) (Fig 1c). This reconfirms that the number of splice variants confers plurality of interactions for proteins to be hubs whereas such a need is not required for proteins to be pure bottlenecks.

### Pathway centrality and evolutionary rates

Evolutionary rates of proteins are related to their expression breadth, expression patterns, essentiality and network topology.^50,51^ Previous studies have shown that degree and betweenness are evolutionarily conserved.^8,9^ It is also reported that hub-non-bottlenecks evolve significantly more slowly than non-hub-bottlenecks.^9^ We investigated the evolutionary rates for three categories of proteins and found that MX proteins evolve at much slower rates than PHs and PBs (P value < 2.2 × 10^-16^) (Fig.2a). Among PHs and PBs, PHs evolve at slower rates which is in agreement with above mentioned previous studies. Hence, proteins with high degree as well as high betweenness are evolutionarily more conserved than the proteins with only high degree and the proteins with only high betweenness values. Again, we found MX proteins involved in multiple pathways (P value < 2.2 × 10^-16^) than PHs and PBs (Fig.2b) which corroborates with their high evolutionary conservation.

**Figure 2:**
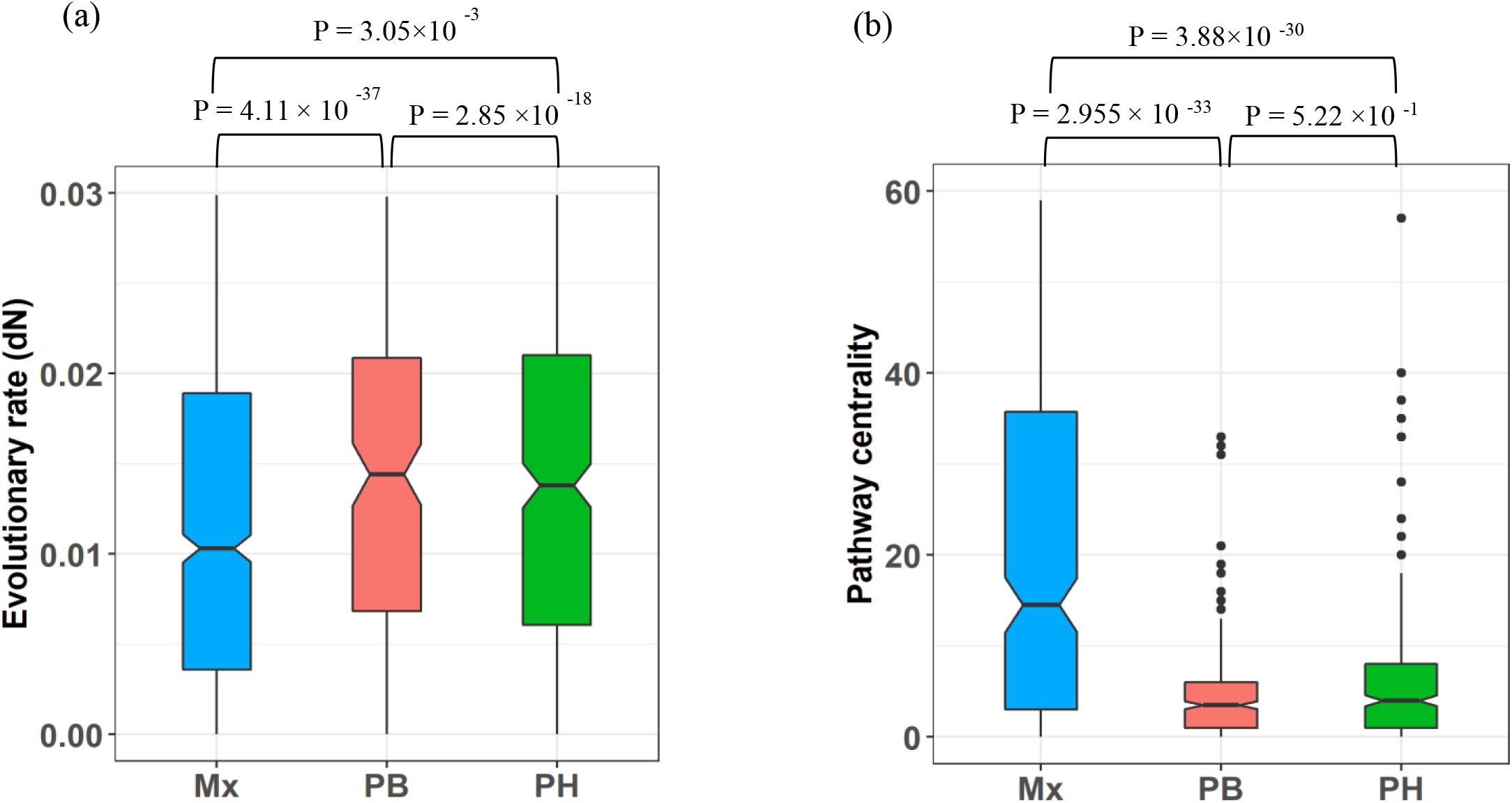
Box plots illustrating (a) Evolutionary rates, (b) Pathway Centrality values for PHs (pure hubs), PBs (pure bottlenecks) and MXs (Mix proteins). It can be seen that MXs show lower evolutionary rates than PBs and PHs (P value < 2.2 × 10^-16^) and involved in a greater number of pathways (P value < 2.2 × 10^-16^) than PHs and PBs. The P-value was calculated by Kruskal-Wallis test.

### Functional enrichment analysis

From previous reports ^47^ it has been shown that kinases, receptors and secreted proteins form hubs in protein-protein interaction network. We also find that these functions are significantly enriched in pure hubs (Fig.3b). Our GO enrichment analysis regarding biological processes reveals that MX proteins are mainly involved in protein metabolism, cell growth and maintenance, cell communication, while PBs are mainly involved in immune response, transport and metabolism. For cellular components, MX proteins are enriched with GO terms related to cytoplasm, nucleus, centrosome, cytosol and centrosome, while PBs are enriched with plasma membrane, extracellular region, lysosome and endoplasmic reticulum. When considering molecular function, we find that MX proteins are primarily enriched in RNA binding, transcription regulator activity, ubiquitin-specific protease activity and receptor signaling complex scaffold activity, whereas PBs are enriched in transporter activity, receptor binding and transmembrane receptor activity. These findings indicate that PHs are involved in core cellular machinery such as transcription and replication as it is part of a single module whereas PBs are involved in infection and signal transduction such as GPCR ligand binding (Fig.3a) and MX proteins are mainly involved in core signaling pathway which requires interconnection between different modules (Fig.3c).

**Figure 3:**
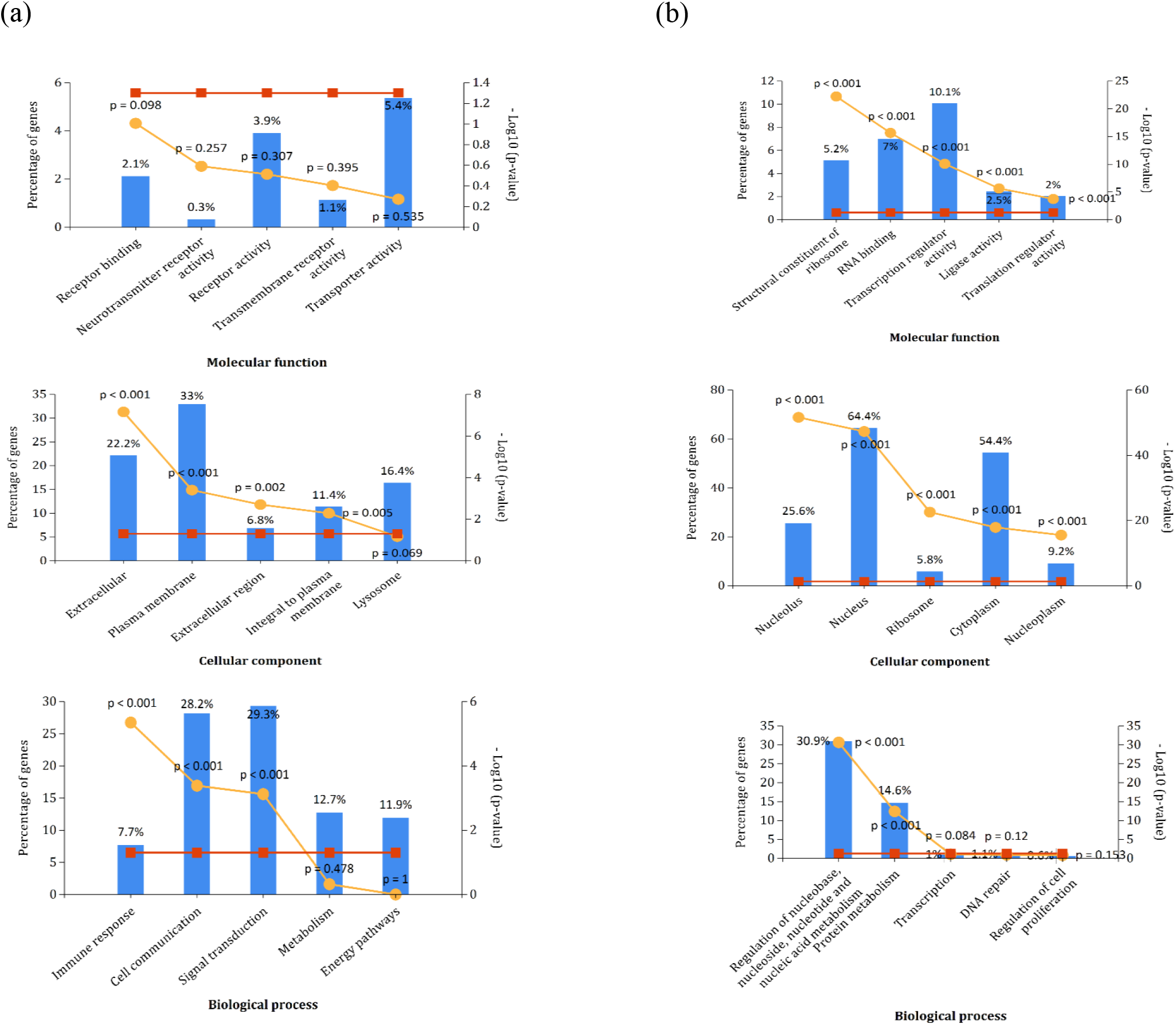
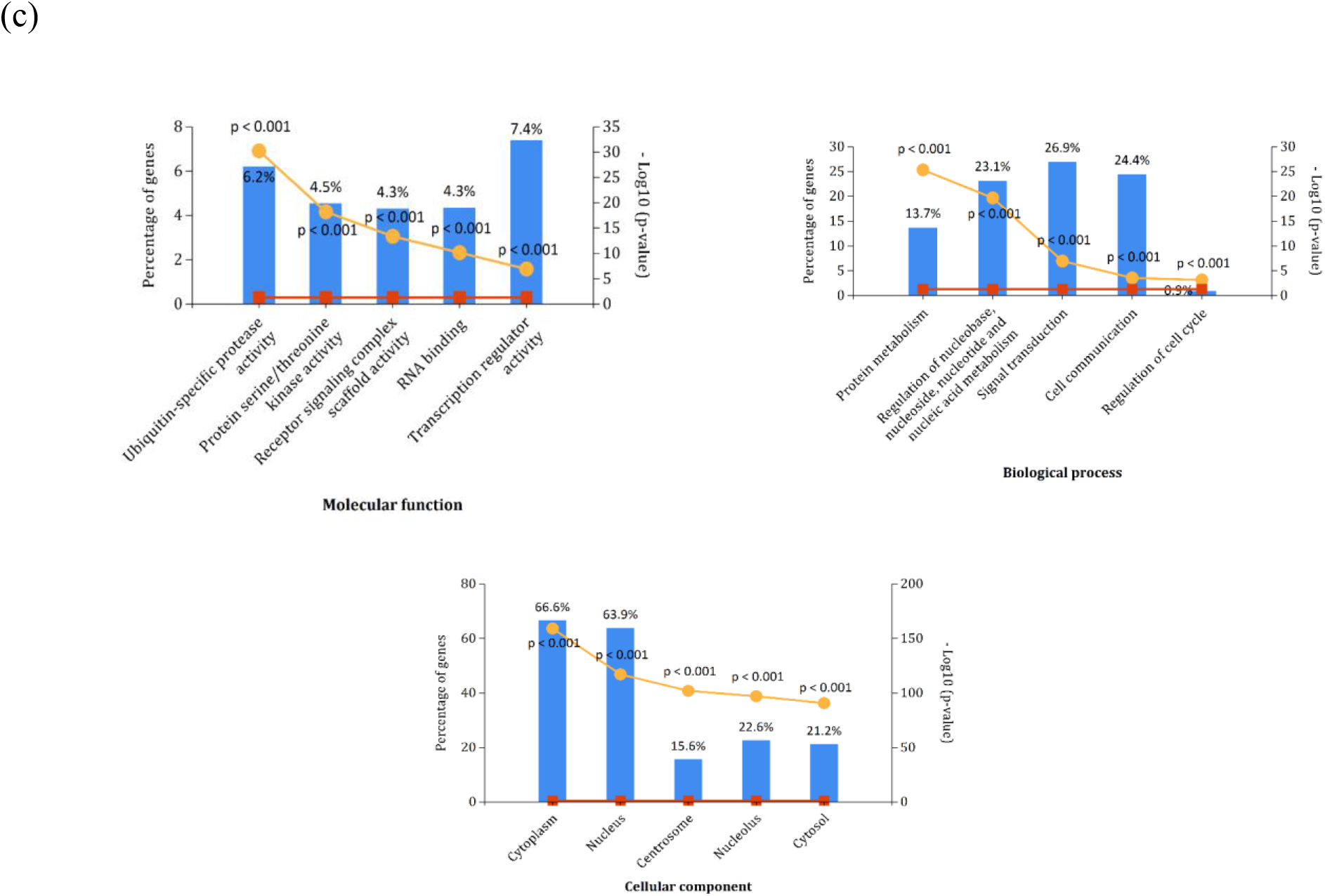
Gene Ontology enrichment analysis for (a) PBs (Pure bottlenecks), (b) PHs (Pure hubs) and (c) MXs (Mix proteins). Blue bars represent the percentage of genes; red line is the reference p value (0.05); yellow line is the relative p value and is significant if less than 0.05.

### Essentiality of PHs, PBs and MXs

Essential proteins play important roles in cellular processes such as development and survival. Among the 926 PHs about 73% are essential proteins, whereas among 2124 MX and 629 PBs the essential proteins form about 23% and 47% respectively. In other words, the products of essential genes are enriched in pure hubs than the other two categories which is in agreement with previous study^4,5,7^ (Table 1). This confirms that essentiality of a protein seems to be solely characterized by its plurality of its interactions in the network.

**Table 1:** The proportion of pure hubs (PHs), pure bottlenecks (PBs) and mix proteins (MX) with different attributes. Z score and the P value are enclosed in the brackets.

| **Attributes** | **Mix Proteins**<br>**(2124)** | **Pure Bottlenecks**<br>**(629)** | **Pure Hubs**<br>**(926)** |
| --- | --- | --- | --- |
| **Pathogen-Targeted**<br>**Proteins (4,585)** | 1328<br>(27.283, $6.74 \times 10^{-164}$) | 183<br>(-0.443, 0.657) | 465<br>(11.184, $4.88 \times 10^{-29}$) |
| **Cancer-related genes**<br>**(1,247)** | 203<br>(2.225, 0.026) | 101<br>(6.912, $4.77 \times 10^{-12}$) | 75<br>(-0.082, 0.934) |
| **Approved Drug**<br>**Targets (1,991)** | 392<br>(6.888, $5.65 \times 10^{-12}$) | 138<br>(6.167, $6.95 \times 10^{-10}$) | 108<br>(-1.172, 0.241) |
| **Essential genes**<br>**(7,282)** | 1541<br>(16.543, $1.79 \times 10^{-61}$) | 295<br>(-0.307, 0.758) | 678<br>(11.216, $3.40 \times 10^{-29}$) |
| **Disease-related**<br>**genes**<br>**(1,716)** | 337<br>(6.340, $2.29 \times 10^{-10}$) | 98<br>(3.236, 0.0012) | 105<br>(0.079, 0.937) |

### Composition of pathogen targeted proteins in PHs, PBs and MXs

Pathogens such as Bacteria, Fungi, Virus and protozoa communicate with human cells through physical interactions with various human proteins. Previous studies have reported that human proteins interacting with bacterial and viral pathogens tend to have higher degrees and high betweenness centrality values in the human PPI network^12^. Investigation of the proteins in the GIN with PHISTO revealed that 4,585 in GIN correspond to those targeted by pathogens. Of these pathogens targeted human proteins, about 50% form PHs, 63% form MXs and 29% form PBs in our GIN. Thus, our analysis reveals that the proteins targeted by pathogens are more often MX proteins than the PHs and PBs (Table 1). It may be worthwhile to recall an earlier report where it was found that the proteins with high betweenness are depleted in bacterial or viral targets.^10^

### Cancer genes and drug targets

In protein-protein interaction networks, hubs and bottlenecks may constitute an important source of drug targets. Therefore, the proteins in the GIN were searched against the DrugBank^40^ (http://www.drugbank.ca/) and found 1,991 drug targets as part of the GIN. We also found that about 12% of PHs, 18% of MXs and 22% of PBs form drug targets. Pure bottlenecks are more enriched in drug targets (Table 1) than the other two categories. It is known that most of the drug targets are receptor proteins such as G-protein coupled receptors and transport proteins. Our analysis of GO functional terms for PBs also had revealed that these proteins are involved in transporter and receptor activities.

We further investigated abundance of disease associated proteins among the three categories. Among PH, PB and MX categories, 11%, 10% and 16% of proteins could be associated to OMIM diseases. This indicated that the MXs i.e., the proteins with higher degree and betweenness are more often likely related to be associated to diseases than PHs and PBs.

We also searched for protein products of the cancer associated genes in the GIN. A total of 1,247 cancer associated proteins were found in the GIN. Furthermore, we also found that the cancer related genes are significantly more abundant in the PBs than MX and PHs (Table 1) and hence corroborates with the previous studies where bottleneck proteins were shown to be mostly comprised of breast and prostate cancer genes.^13,15^

### Proximity analysis of PHs, PBs and MXs to the drug targets

We analyzed the path length between the known approved drug targets and the different categories of proteins. For every pathlength value (0 means the protein is the drug target, 1 means the pathlength between a drug target and the protein of interest is 1, etc) we calculated the proportion of PHs, PBs and MXs and the results are shown in Fig. 4. From this figure it can be seen that the drug targets are more frequently comprised of proteins other than PHs, PBs and MXs. However, among the three categories, MXs form more often the drug targets than PHs and PBs and this pattern can be seen even with the proteins at pathlengths up to 4 and at longer pathlengths only PBs are found. This analysis establishes the role of proteins with high betweenness values not themselves as direct drug targets but as mediators to spread the effect to the complete network.

**Figure 4:**
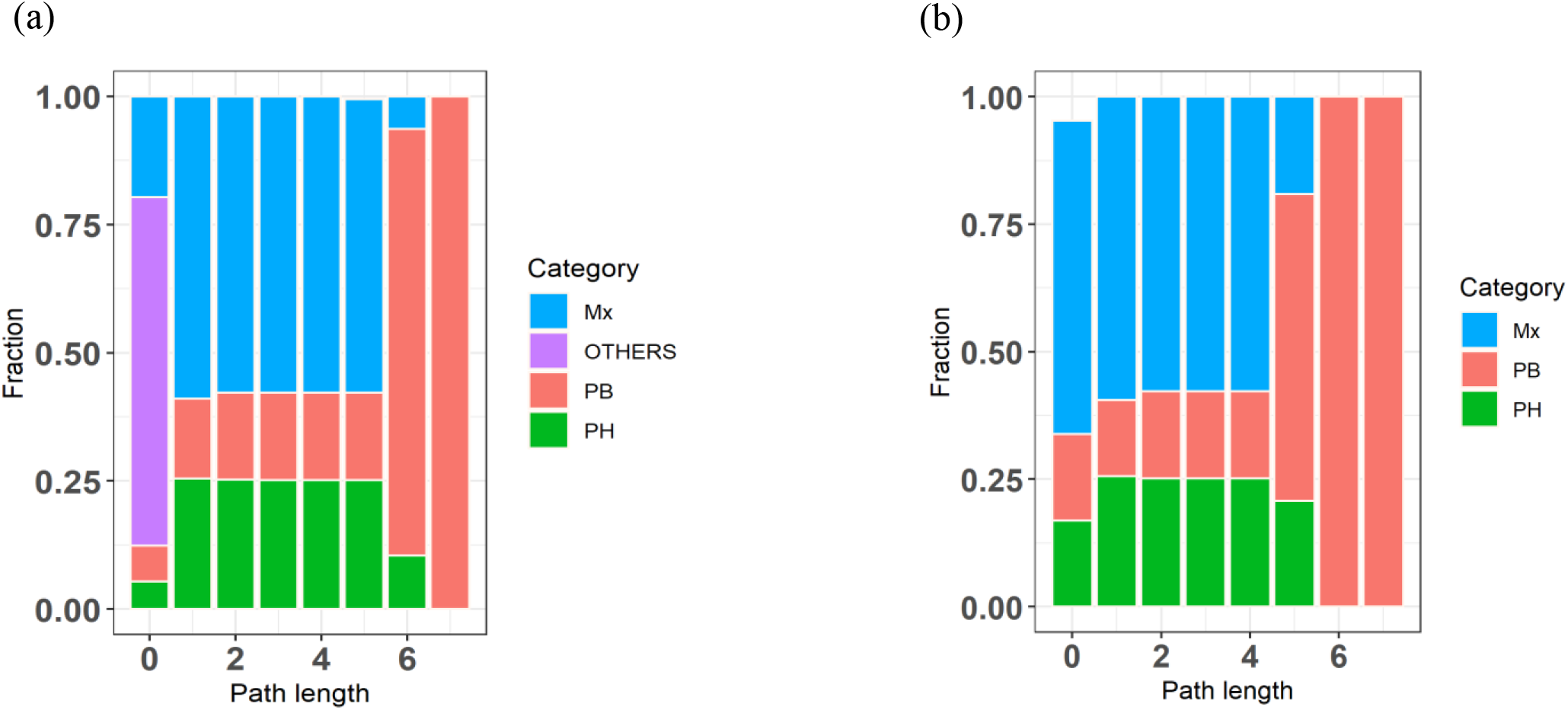
The path length between approved drug targets and PH (Pure hubs), PB (Pure bottlenecks), MX (Mix proteins) and others (proteins other than PHs, PBs and MXs in the GIN).

## Discussion and Conclusion

In this study, we have identified three groups of proteins and find surprising outcomes concerning the importance of making distinction between pure hubs (PHs), pure bottlenecks (PBs) and those with high degree and betweenness centralities (MXs).

PHs are mostly disordered proteins with large number of splice variants enriched with binding sites in their disordered regions. These structural features seem to support PHs functional roles in transcription, replication, DNA repair and translation etc., which are very essential for the survivability of cells. Thus, this study reconfirms that degree is a better predictor of essentiality than betweenness in interaction networks. An example of a PH is Cyclin Dependent Kinase13 (CDK13, UniprotKB: Q14004), which belongs to a member of cyclin-dependent serine/threonine protein kinase family. CDK13 has an essential role in cell cycle control. This protein plays an important role in regulation of haematopoiesis and mRNA processing. In HIV-1 infection, the splicing is regulated by viral protein Tat. CDK13 interacts with Tat protein and increases HIV-1 mRNA splicing.^52^ But, the silencing of CDK13 will increase the HIV-1 production. CDK13 might be the possible essential restriction factor for viral replication.

The fact that MXs are involved in large number of KEGG pathways and that they evolve slowly seem to correlate with each other. These proteins as well as PHs are often targeted by pathogens but not PBs suggest that pathogens target only the highly connected proteins in the network but not with high betweenness which is contrary to previous study.^10^

Our studies suggest that PBs form the drug targets as well as associated with cancer genes and are involved in receptor mediated signaling complex process, transport and receptor activity. For example, Bone Morphogenetic protein 4 (BMP4, UniprotKB: P12644) is a pure bottleneck. It is an extracellular multi-signaling molecule which encodes a secreted ligand of Transforming Growth Factor - Beta (TGF-beta). It is involved in the transport and activation of the SMAD family of proteins and in the development of many organ systems, as well as adult tissue homeostasis. Mutations in BMP4 gene are associated with several human diseases like cancers, multiple cardiovascular diseases,^53^ orofacial cleft and microphthalmia.^54^ and hence this protein has been suggested as a possible therapeutic target in cancer cells.^55^

In summary, this study unravels some distinct characteristic features associated with pure hubs, pure bottlenecks as well as the proteins having both high degree and high betweenness centralities. Studies reported elsewhere had majorly focused on properties of hubs ignoring their possible high betweenness values or properties of bottlenecks ignoring their possible high degrees. The present study has facilitated clear understanding of distinct attributes of the above mentioned three distinct centrally important proteins. The distinct structural and functional characteristics associated with each of these categories would help in prioritizing network-based target selections toward developing novel drugs as well as developing novel therapeutic regimes for communicable as well as non-communicable diseases. Similar analysis can be extended to other biological networks as a means to understand biological systems in-depth.

## FIGURES

**Figure S1:**
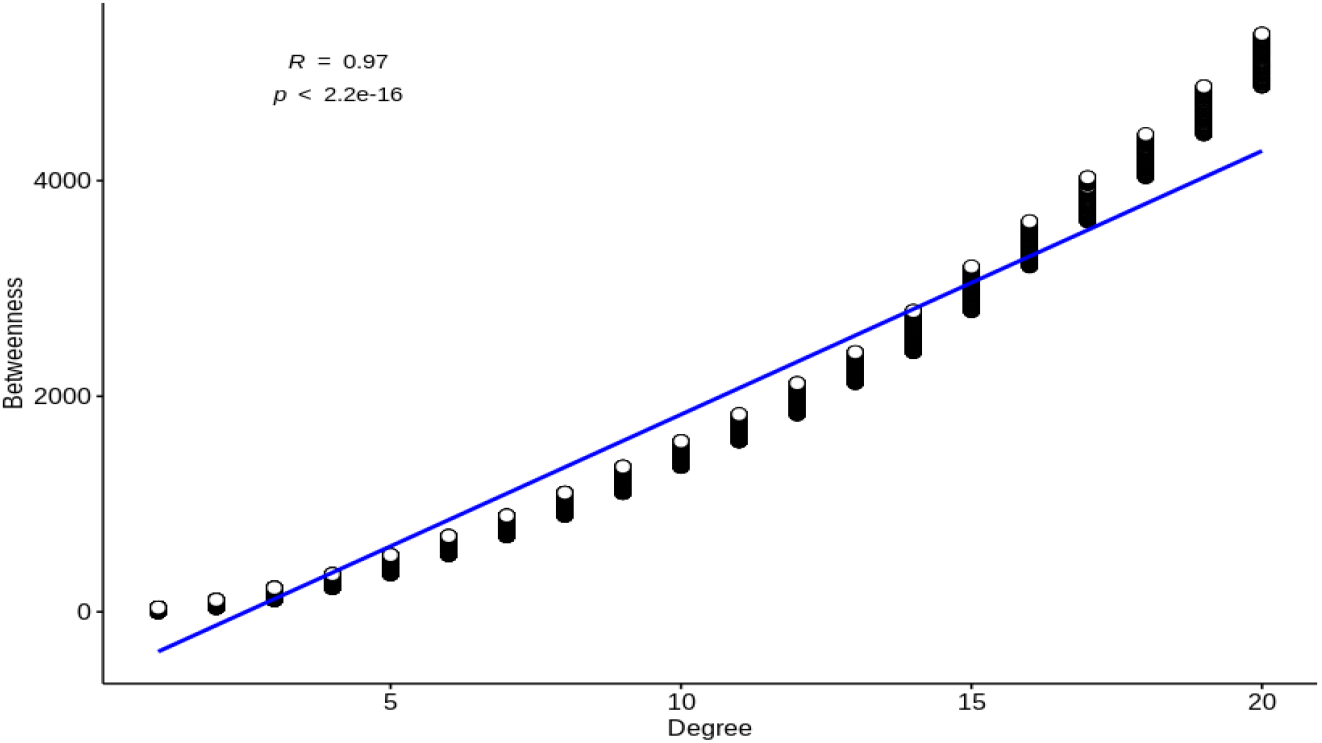
Plot representing significant positive correlation between degree and betweenness (Pearson correlation coefficient of 0.97, P value < 2.2×10 ^-16^)

**Figure S2:**
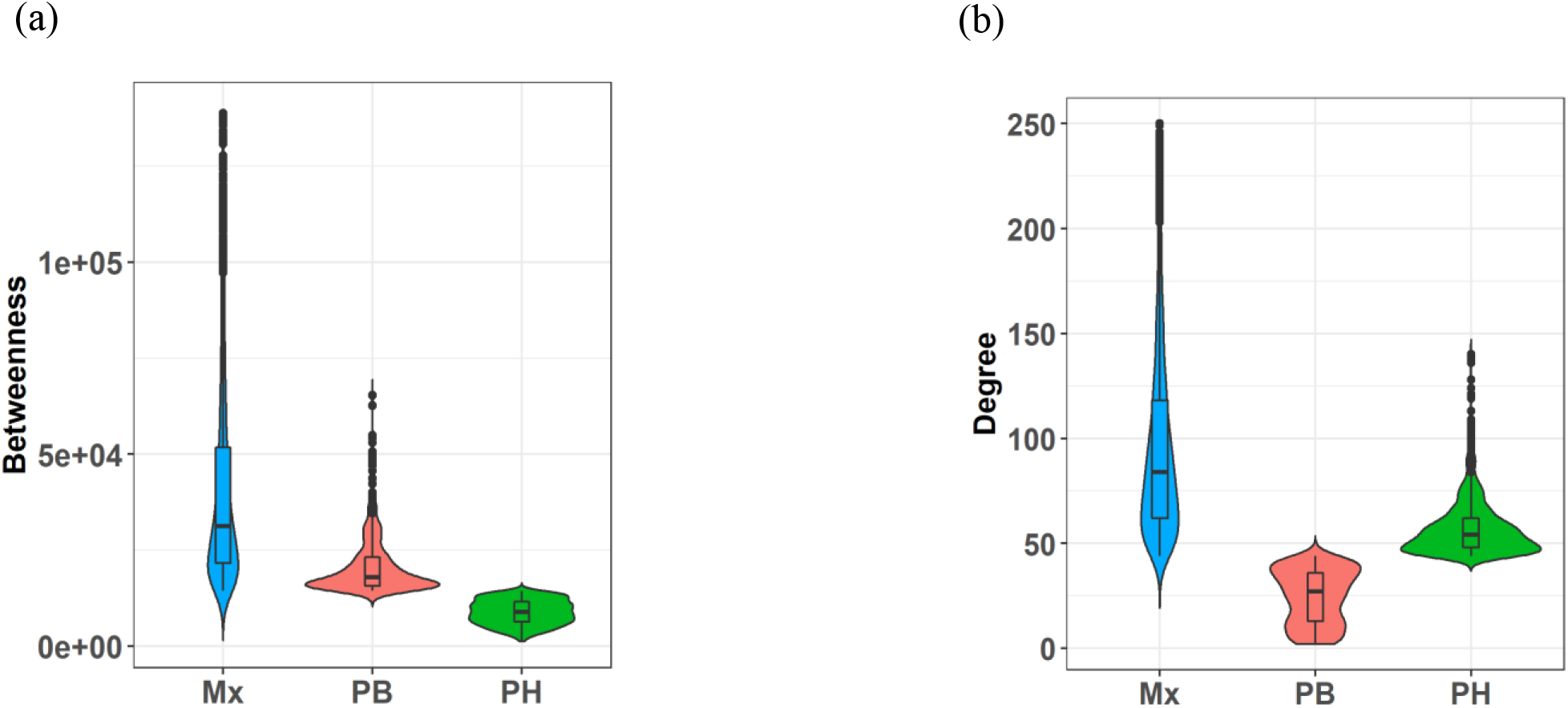
Violin plot of (a) Betweenness distribution, (b) Degree distribution. Mix proteins (Mx) have higher betweenness values (P value < 1.9×10 ^-14^) and degrees (P value < 2.2×10 ^-16^) as compared with pure hubs (PHs) and pure bottlenecks (PB). P-value was calculated by Kruskal-Wallis test.

## AUTHOR INFORMATION

### Notes

The authors declare no competing financial interest.

## ACKNOWLEDGMENT

N.C., a registered PhD student at University of Hyderabad, gratefully acknowledges the Boarding cum Lodging (BBL) fellowship from University of Hyderabad. HAN and MK gratefully acknowledge the core grant support from the University of Hyderabad. The Bioinformatics Infrastructure Facility (BIF) at the School of Life Sciences is also gratefully acknowledged.

## ABBREVIATIONS

PHs: pure hubs
PBs: pure bottlenecks
MXs: mix proteins
PPINs: protein-protein interaction networks
PPI: protein-protein interaction
HIPPIE: human integrated Protein-Protein interaction rEference
MINT: the molecular iNTeraction database
BioGrid: biological general repository for interaction datasets
DIP: database of interacting proteins
BIND: biomolecular interaction network database
HPRD: human protein reference database
MIPS: munich information center for protein sequences
KEGG: kyoto encyclopedia of genes and genomes
GO: gene ontology
MF: molecular function
BP: biological process
CC: cellular component
DEG: database of essential genes
PHISTO: pathogen-host interaction search tool
STRING: search tool for the retrieval of interacting genes/proteins
APID: agile protein interaction data analyzer
iRefIndex: interaction reference index
MPIDB: Microbial Protein Interaction Database
OMIM: online mendelian inheritance in man
GWAS: genome-wide association studies
GIN: global interaction network
IDPs: intrinsically disordered proteins
CDK13: cyclin dependent kinase13

## REFERENCES

1. Basi, A. L., Oltvai, Z.N. Network biology: understanding the cell’s functional organization. Nat Rev Genet. 2004, 5(2), 101–13.

2. de Juan, D., Pazos, F., Valencia, A. Emerging methods in protein co-evolution. Nat Rev Genet. 2013, 14(4), 249–61.

3. Vidal, M., Cusick, M. E., Barabasi, A. -L. Interactome networks and human disease. Cell. 2011,144(6), 986–98.

4. Jeong, H., Mason, S. P., Barabási, A. -L., Oltvai, Z.N. Lethality and centrality in protein networks. Nature. 2001, 411, 41–42.

5. Yu, H., Kim, P. M., Sprecher, E., Trifonov, V., Gerstein, M. The Importance of Bottlenecks in Protein Networks: Correlation with Gene Essentiality and Expression Dynamics. PLoS Comput Biol. 2007, 3(4), e59.

6. Girvan, M., Newman, M. E. Community structure in social and biological networks. Proc Natl Acad Sci U S A. 2002, 99, 7821–7826.

7. Kim, P. M., Korbel, J. O., Gerstein, M.B. Positive selection at the protein network periphery: evaluation in terms of structural constraints and cellular context. Proc Natl Acad Sci U S A. 2007, 104(51), 20274–20279.

8. Pang, E., Hao, Y., Sun, Y. Differential variation patterns between hubs and bottlenecks in human protein-protein interaction networks. BMC Evol Biol. 2016, 260.

9. Joy, M. P., Brock, A., Ingber, D. E., Huang, S. High-betweenness proteins in the yeast protein interaction network. J Biomed Biotechnol. 2005, 96–103.

10. Dyer, M. D., Murali, T. M., Sobral, B.W. The Landscape of Human Proteins Interacting with Viruses and Other Pathogens. PLoS Pathog. 2008, 4(2), e32.

11. Calderwood, M. A., Venkatesan, K., Xing, L., Chase, M. R., Vazquez, A., Holthaus, A. M., Ewence, A. E., Li, N., Hirozane-Kishikawa, T., Hill, D. E., Vidal, M., Kieff, E., Johannsen. Epstein–Barr virus and virus human protein interaction maps. Proc. Natl. Acad. Sci. U.S.A. 2007, 104, 7606–7611.

12. Schleker, S., Trilling M. Data-warehousing of protein-protein interactions indicates that pathogens preferentially target hub and bottleneck proteins. Front Microbiol. 2013, 4.

13. Ozgür, A., Vu, T., Erkan, G., Radev, D.R. Identifying gene-disease associations using centrality on a literature mined gene-interaction network. Bioinformatics. 2008, 24(13), i277–85.

14. Goñi, J., Esteban, F. J., de Mendizábal, N., Sepulcre, J., Trevijano, S. A., Agirrezabal, I., Villoslada, P. A computational analysis of protein-protein interaction networks in neurodegenerative diseases. BMC Syst Biol. 2008, 2(1), 52.

15. Bafna, D., Isaac, AE., Identification of Target Genes in Breast Cancer Pathway using Protein-Protein Interaction Network. International Journal of Cancer research. 2017, 13(2), 51–58.

16. Feng, Y., Wang, Q., Wang, T., Drug Target Protein-Protein Interaction Networks: A Systematic Perspective. Biomed Res Int. 2017, 1–13.

17. Fu, Y., Guo, Y., Wang, Y., Luo, J., Pu, X., Li, M., Zhang, Z. Exploring the relationship between hub proteins and drug targets based on GO and intrinsic disorder. Computational Biology and Chemistry, 2015, 56, 41– 48.

18. Yao, L., Rzhetsky, A., Quantitative systems-level determinants of human genes targeted by successful drugs. Genome Res. 2008, 18(2), 206–213.

19. Goh, K. I., Oh, E., Kahng, B., Kim, D. Betweenness centrality correlation in social networks. Phys Rev E Stat Nonlin Soft Matter Phys, 2003, 67, 017101.

20. Alanis-Lobato, G., Andrade-Navarro, M. A., Schaefer, M.H. HIPPIE v2.0: enhancing meaningfulness and reliability of protein–protein interaction networks. Nucleic Acids Res. 2017, 45.

21. Chatr-aryamontri, A., Ceol, A., Palazzi, L. M., Nardelli, G., Schneider, M. V., Castagnoli, L., Cesareni, G. MINT: the Molecular INTeraction database. Nucleic Acids Res. 2007, 35, D572–D574.

22. Stark, C., Breitkreutz, B. J., Reguly, T., Boucher, L., Breitkreutz, A., Tyers, M. BioGRID: a general repository for interaction datasets. Nucleic Acids Res, 2006,34, D535–D539.

23. Salwinski, L., Miller, C. S., Smith, A. J., Pettit, F. K., Bowie, J. U., Eisenberg, D. The database of interacting proteins: 2004 update. Nucleic Acids Res. 2004, 32, D449–D45.

24. Isserlin, R., El-Badrawi, R.A., Bader, G.D. The biomolecular interaction network database in PSI-MI 2.5. Database (Oxford) 2011, baq037.

25. Kerrien, S., Aranda, B., Breuza, L., Bridge, A., Broackes-Carter, F., Chen, C., Duesbury, M., Dumousseau, M., Feuermann, M., Hinz, U., Jandrasits, C., Jimenez, R., Khadake, J.; Mahadevan, U., Masson, P., Pedruzzi, I., Pfeiffenberger, E., Porras, P., Raghunath, A., Roechert, B., Orchard, S., Hermjakob, H. The IntAct molecular interaction database in 2012. Nucleic Acids Res. 2012, 40, D841–D846.

26. Keshava Prasad, T. S., Goel, R., Kandasamy, K., Keerthikumar, S., Kumar, S., Mathivanan, S., Telikicherla, D., Raju, R., Shafreen, B., Venugopal, A., Balakrishnan, L., Marimuthu, A., Banerjee, S., Somanathi, D. S., Sebastian, A., Ray, S., Kishore, C.J.H., Kanth, S., Ahmed, M., Kashyap, K. M., Mohmood, R., Ramachandra, Y. L., Krishna, V., Rahiman, A. B., Mohan, S., Ranganathan, P., Ramabadran, S., Chaerkady, R., Pandey, A. Human protein reference database–2009 update. Nucleic Acids Res. 2009, 37, D767–D772.

27. Pagel, P., Kovac, S., Oesterheld, M., Brauner, B., Dunger-Kaltenbach, I., Frishman, G., Montrone, C., Mark, P., Stumpflen, V., Mewes, H. W., Ruepp, A., Frishman, D. The MIPS mammalian protein-protein interaction database. Bioinformatics. 2005, 21, 832–834.

28. Mészáros, B., Erdős, G., Dosztányi, Z. IUPred2A: context-dependent prediction of protein disorder as a function of redox state and protein binding. Nucleic Acids Res. 2018, 46(W1), W329–W337.

29. Meszaros, B., Simon, I., Dosztanyi, Z. Prediction of protein binding regions in disordered proteins. PLoS Comput. Biol. 2009, 5 (5), e1000376.

30. Kanehisa, M., Goto, S., KEGG: Kyoto encyclopedia of genes and genomes. Nucleic Acids Res. 2000, 28, 27– 30.

31. Hubbard, T., Barker, D., Birney, E., Cameron, G., Chen, Y,; Clark, L., Cox, T., Cuff, J., Curwen, V., Down, T., Durbin, R., Eyras, E., Filbert, J., Hammond, M., Huminiecki, L., Kasprzyk, A., Lehvaslaiho, H., Lijnzaad, P., Melsopp, C., Mongin, E., Pettett, R., Pocock, M., Potter, S,; Rust. A., Schmidt. E., Searle, S., Slater, G., Smith, J., Spooner, W., Stabenau, A., Stalker, J., Stupka, E.,Ureta-Vidal Vastrik. I., Clamp, M. The Ensembl genome database project. Nucleic Acids Res. 2002, 30, 38–41.

32. Kinsella, R. J., Kahari, R. J., Haider, S., Zamora, J., Proctor, G., Spudich, G., King, J., Staines, D., Derwent, P., Kerhornou, A., Kersey, P., Flicek, P. Ensembl BioMarts: a hub for data retrieval across taxonomic space. Database (Oxford), 2011.

33. Pathan, M., Keerthikumar, S., Ang. C.S., Gangoda, L., Quek, C.Y.J., Williamson, N. A., Mouradov, D., Sieber, O. M., Simpson, R. J., Salim, A., Bacic, A., Hill, A. F., Stroud, D.A., Ryan, M.T., Agbinya, J. I., Mariadason, J. M., Burges, A.W., Mathivanan, S. FunRich: An open access standalone functional enrichment and interaction network analysis tool. Proteomics. 2015;15(15), 2597–2601.

34. Zhang, R., Lin, Y. Deg 5.0, a database of essential genes in both prokaryotes and eukaryotes. Nucleic Acids Res. 2009, 37.

35. Szklarczyk, D., Morris, J. H., Cook, H., Kuhn, M., Wyder, S., Simonovic, M., Santos, A., Doncheva, N. T., Roth, A., Bork, P., Jensen, L. J., Mering, C. V. The STRING database in 2017: Quality-controlled protein-protein association networks, made broadly accessible. Nucleic Acids Res. 2017, 45, D362–D368.

36. Prieto, C., De Las Rivas, J. APID: Agile Protein Interaction DataAnalyzer, Nucleic Acids Res., 2006, 34, W298–W302.

37. Croft, D., Mundo, A. F., Haw, R., Milacic, M., Weiser, J., Wu G, et al. The Reactome pathway knowledgebase. Nucleic Acids Res. 2014, 42(Database issue), D472–7.

38. Razick, S., Magklaras, G., Donaldson, I.M. iRefIndex: a consolidated protein interaction database with provenance. BMC Bioinformatics, 2008, 9, 405.

39. Goll, J., Rajagopala, S. V., Shiau, S. C., Wu, H., Lamb, B. T., Uetz, P. MPIDB: the microbial protein interaction database, Bioinformatics, 2008, 24, 1743–1744.

40. Wishart, D. S., Knox, C., Guo, A. C., Cheng, D., Shrivastava, S., Tzur, D., Gautam, B., Hassanali, M. DrugBank: a knowledgebase for drugs, drug actions and drug targets. Nucleic Acids Res, 2008, 36, D901–D906.

41. Amberger, J., Bocchini, C. A., Scott, A. F., Hamosh, A, McKusick’s. Online Mendelian Inheritance in Man (OMIM), Nucleic Acids Res. 2009, 37, D793–D796.

42. Welter, D., MacArthur, J., Morales, J., Burdett, T., Hall, P., Junkins, H., Klemm, A., Flicek, P., Manolio, T., Hindorff, L., Parkinson, H. The nhgri gwas catalog, a curated resource of snp-trait associations. Nucleic Acids Res. 2014, 42(D1), 1001–6.

43. Dessau, R. B., Pipper, C.B. [″R″–project for statistical computing]. Ugeskr. Laeg. 2008, 170 (5), 328−30.

44. Wright, P.E., Dyson, H.J. Intrinsically unstructured proteins: Re-assessing the protein structure–function paradigm. J Mol Biol. 1999, 293: 321–331.

45. Dunker, A.K., Obradovic, Z. The protein trinity—Linking function and disorder. Nat Biotechnol. 2001,19: 805–806.

46. Haynes, C., Oldfield, C. J., Ji, F., Klitgord, N., Cusick, M. E., Radivojac, P., Uversky, V. N., Vidal, M., Iakoucheva, L.M. Intrinsic disorder is a common feature of hub proteins from four eukaryotic interactomes. PLoS Comput. Biol. 2006, 2 (8), e100.

47. Kiran, M., Nagarajaram, H.A. Global versus local hubs in human protein-protein interaction network. J Proteome Res. 2013, 12(12), 5436–5446.

48. Sinha, A., Nagarajaram, H.A. Effect of alternative splicing on the degree centrality of nodes in protein-protein interaction networks of Homo sapiens. J Proteome Res. 2013, 12(4), 1980–1988.

49. Yura, K., Shionyu, M., Hagino, K., Hijikata, A., Hirashima, Y., Nakahara, T., Eguchi, T., Shinoda, K., Yamaguchi, A., Takahashi, K., Itoh, T., Imanishi, T., Gojobori, T., Go, M,; Alternative splicing in human transcriptome: functional and structural influence on proteins. Gene. 2006, 380(2), 63–7.

50. Pal, C., Papp, B., Lercher, M.J. An integrated view of protein evolution. Nat Rev Genet. 2006, 7(5), 337–48.

51. Zhang, J., Yang, J-R. Determinants of the rate of protein sequence evolution. Nat Rev Genet. 2015, 16(7), 409–20.

52. Berro, R., Pedati, C., Kehn-Hall, K., Wu, W., Klase, Z., Even, Y., Guinevere, A-M., Ammosova, T., Nekhai, S., Kashnchi, F. CDK13, a New Potential Human Immunodeficiency Virus Type 1 Inhibitory Factor Regulating Viral mRNA Splicing. J Virol. 2008.

53. Morrell, N. W., Bloach, D. B., Dijke, P. T., Goumans, M-J. T.H., Hata, A., Smith, J., Yu, P. B., Bloch, K. D. Targeting BMP signalling in cardiovascular disease and anaemia. Nat. Rev. Cardiol. 2016, 13, 106–120.

54. Zhang, X., Li, S., Xiao, X., Jia, X., Wang, P., Shen, H., Guo, X., Zhang, Q. Mutational screening of 10 genes in Chinese patients with microphthalmia and/or coloboma. Mol Vis 2009, 15, 2911–2918.

55. Li, B. Bone Morphogenetic Protein-Smad Pathway as Drug Targets for Osteoporosis and Cancer Therapy. Endocrine, Metab Immune Disord Targets. 2008, 8(3), 208–219.

